# Circular RNAs may embed pieces of real-world sensory information into an episodic memory

**DOI:** 10.1101/2022.10.06.511043

**Authors:** Arun Asok

## Abstract

For a generation, neuroscience has searched for a molecule that stores our memories across time. This search has focused on proteomic mechanisms, but less is known about RNA. Here, we identify a new persisting class of RNA associated with long-term memory – Circular RNAs. Unlike other RNAs, Circular RNAs are stable for days or longer and may provide a means for storing sensory information across time. We leveraged a differential fear conditioning paradigm whereby individual mice sample all real-world sensory inputs (i.e., auditory, visual, gustatory, olfactory, and incidental tactile) in a quasi-stochastic manner prior to receiving different intensities of an unconditioned stimulus (US) foot-shock. While Pavlovian models of learning from the 20^th^ century were critical for understanding elemental associations, they fail to appreciate (1) what US content remains inside of a complex conditioned stimulus (CS) or response (CR – a behavioral manifestation of an episodic memory), (2) what happens when the associations involve multiple senses, and (3) what biologically happens to the real-world US. Given (1) we are constantly sampling information from our environment through all our senses and (2) the US at a given moment in time likely adds value to imprint that multisensory representation, we propose the real-world US is biologically encoded via back-spliced Circular RNAs within the cells and circuits that represent a particular episodic memory and present days later. This logic, best simplified by the equation: 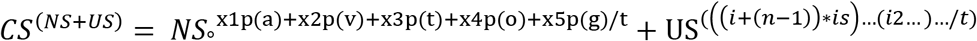, allowed us to ask how the formation of similar episodic memories, which only differ in relation to the content of US information, alter Circular RNAs in the CA1 subfield of the hippocampus – a brain area critical for episodic memories. We found that stronger foot-shock USs during conditioning produce stronger memories relative to weaker USs 24-h later. Stronger memories also generalize to novel/safe environments 48-h later. Moreover, the unconditioned response is highly correlated with future CRs, suggesting (1) an understudied relationship between the strength and type of US/URs and future CRs in complex environments as well as (2) fear generalization, at least in the short-term, is associated with the embedding of additional US information. Next-generation Circular RNA sequencing 1-hr after acquisition revealed a remarkably small set of circular RNAs relative to nearly identical, yet weaker, episodic memories in CA1. Gene Ontologies for mice that formed weaker and stronger memories matched those families classically involved in weaker and stronger forms of memory across species. Preliminary *in situ* hybridization visually confirmed the presence of Circular RNAs in the CA1 subfield. Future experiments will examine the persistence of Circular RNAs in cells of a memory trace (i.e., engram cells; in situ hybridization) at recent (4 days) and remote (21-days) time points. Taken together with our mathematical model for multisensory learning, our data suggest that Circular RNAs do not contribute to the storage of the multisensory configural representation, but perhaps to the storage of discrete pieces of real-world *sensory* information related to the US that is partially embedded inside of a memory trace early-on. Importantly, in the above model for multisensory learning, the discrete USs are biologically separable from the future *CS^(NS+US)^* associations and US strength is modifiable across time. This work reveals fundamental insights into how we store pieces of real-world sensory information in an episodic memory at the biological level of the brain.

**One Sentence Summary:** circRNAs biologically encode real-world sensory information into a long-term memory

## Introduction

We are who we are in part by what we learn, but more importantly by what we remember (*1–3*). The experiences of our lives are stored as “episodes” that represent a synthesis of the sensory stimuli acquired from our various senses (e.g., olfactory, gustatory, auditory, visual, and, critically, tactile; cf. (*4–7*)). Episodic memories are essential for adaptive behavior and are also the most notably affected in anxiety and post-traumatic stress disorders (PTSD; (*8–14*)). Despite considerable progress in identifying how sensory experiences from the real-world are associated with precise biological changes (e.g., ex vivo, in vivo, and at the level of neuronal networks (*7, 15–22*)), less is known about which molecules within neuronal networks store information at the biological level.

The search for a lasting biological mark of memory dates back nearly half a century, to the early 1980s where Francis Crick outlined ideas for an intramolecular autocatalytic enzyme capable of encoding information to allow a memory to persist across time (*23*). Indeed, empirical support for the above ideas began to emerge in the mid to late 1980s with the identification of CAMKII autophosphorylation (*24*) as well as discoveries in the 1990s (*25, 26*) and early 2000s on an atypical, constitutively active Protein-Kinase C isoform, PKMζ (*27–29*).

These early studies were paralleled by groundbreaking work on pathogenic prions in the 1980s by Stanley Prusiner. Prusiner’s group identified several biological properties of prions which alluded to a role as a stable, lasting biological information storage device. Subsequent studies (*30*) by Eric Kandel and colleagues leveraged Prusiner’s insight into pathogenic prions to discover functional prions in the brain (*31, 32*). Functional prions are different from their pathogenic counterparts in that they contain Q/N rich ‘prion-like’ domains and aggregate in an activity dependent manner but are non-pathogenic. Importantly, functional prions are also RNA-binding proteins which make them special relative to other proteins given they can bind various classes of RNA (*30–35*). Moreover, recent work has shown that well-known genes related to synaptic plasticity have a unique capacity to bind RNA and move between cells – suggesting a more complex role for RNA in memory than previously thought (*36, 37*).

In recent decades, work from several laboratories revealed how RNA-binding proteins such as T-cell intracellular antigen 1 (TIA-1) or cytoplasmic polyadenylation binding protein 3 (CPEB3) may contribute to the persistence, and perhaps pathologies, of memory (cf. (*31*)). Moreover, we’ve made considerable progress in identifying how a signal moves from the presynaptic to postsynaptic neuron to influence semi-linear transcription and the translation of proteins necessary for synaptic plasticity and ultimately long-term memory (e.g., CAMKII, PKAs, MAPKs, and CREB, neurotrophins, RNA-binding proteins, and other autocatalytic enzymes (*25, 26, 38–40*)). However, the challenging question remains: How are the sensory components of a real-world experience biologically stored in cells, incorporated into an episodic memory, and physically represented across time?

Studies continue to search for an intramolecular autocatalytic enzyme associated with long-term memory. The majority of these studies have logically focused on proteins given they are the functional “end”-product of mRNA translation and have a diverse array of longer-lasting functions (*41*). However, proteins generally have short half-lives functioning on the time scale of hours to a few days (*42, 43*)). In recent years, research has shifted towards identifying how RNA, rather than proteins, may serve as a mechanism for encoding information at the biological level (*6, 44*). Indeed, Crick and others in the latter half of the 20^th^ century hypothesized that RNA may provide a mechanism for carrying information across generations (*23, 45, 46*). Curious studies littered throughout the fringes of the past half-century, and recently revisited (*47*), have suggested that RNA may provide a stable platform for encoding information – thereby providing a biological link between our sensory world and biological information encoding (*48, 49*). In fact, work on piRNAs (*50*), long non-coding RNAs (*51*), as well as other RNA species (*49*) have provided compelling evidence for a critical, yet poorly defined role for RNA in memory.

Recently, a new class of RNA was rediscovered – Circular RNA. We know messenger RNA (mRNA) is linearly transcribed from DNA prior to splicing and later translation (e.g., CREB, C/EBP, CPEB3, etc. (*52–55*)). By contrast, circular RNAs are back spliced from exonic and/or intronic regions of DNA. Circular RNAs are unique relative to every other RNA because [1] they are not subject to normal degradation by RNAases, [2] the key RNAase necessary for their degradation is not present in tissue, and [3] they persist for very long time periods (*56, 57*). Studies have recently started to identify circular RNAs in the hippocampus – a brain region critical to episodic memory (*58–61*). However, the function of Circular RNAs remains unknown (*62*). Given (1) their stability on time-scales necessary for episodic memory and (2) their presence in the hippocampus – a key brain region critical for episodic memory – we hypothesized that circular RNAs may provide a stable mechanism (e.g., a real-world information packet) for biologically storing key pieces of unconditioned sensory information inside of an episodic memory.

Research in the past century has used Pavlovian learning principles to understand elementary forms of learning where a neutral stimulus (NS) is paired with an unconditioned stimulus (US) to form a conditioned stimulus (CS) and conditioned response (CR) (*63–66*). This work has provided valuable insights into the associative strength (e.g., ΔV) between single and compound cues and the unconditioned stimulus (US) to produce a future behavioral CR (cf. (*67*)). Moreover, these principles have been essential for understanding how memories are held in discrete, but distributed, networks of cells and circuits (e.g., “the engram”; (*7, 18, 19, 68*). However, Pavlovian models do not mathematically or biologically account for what happens to information about the US after the initial learning, or how US information is stored inside of complex multisensory representations. Given we are continually sampling information from our environment, we hypothesized that USs from the world serve to lock value into a particular multisensory configural representation (CS) within a given moment in time. Moreover, we know that the US is discrete and enters the brain through a specific sensory modality and therefore should be biologically detectable for prolonged timescales if the US is in fact encoded into an associative CS-US memory.

Using the above logic, we reasoned that if we treated sampling of the real-world through the five senses (i.e., auditory, visual, olfactory, gustatory, and incidental tactile) as “static” in time (assuming the sampling of these inputs to stochastically fluctuate as normal) during the formation of an episodic memory, we could precisely increase the tactile US inputs to identify (1) stable circular RNAs associated with a discrete real-world input (i.e., a tactile US) that is (2) located in the major memory center of the brain – the hippocampus, and (3) perhaps incorporated into the cells of a memory trace (i.e., engram cells). Our findings provide the first insight into how a novel class of RNA may provide a stable, activity-dependent, biological storage mark for real-world unconditioned sensory information that is embedded into a memory trace and operates on timescales akin to human memory storage. Moreover, our framework provides a dual-process biological theory whereby the CS is encoded via forward spliced mRNA transcription and the US is encoded via the back spliced Circular RNAs to form a complex multisensory episodic memory.

## Results

We hypothesized that circular RNAs (circRNAs) may serve as a link between the sensory world and the biological deposition of unconditioned stimulus information into an episodic memory. Thus, we first sought to examine if we could finely control one sensory input during the formation of an episodic memory in mice. Given we cannot fully predict all of the sensory stimuli an animal is sampling at a given moment in time (*22, 69, 70*), we allowed for sampling of the sensory environment (i.e., the “multisensory context”) to stochastically vary as normal and treated this as a static variable (described below). We focused on manipulating a single sensory input similar to past work (*5, 71*) and (*72*). We synthesized a half-century’s worth of findings on unconditioned stimulus (US) parameters, unconditioned response (UR) defensive behaviors (*73–75*), contextual learning (*8, 71, 76, 77*), and the molecular biology of long-term memory storage (cf. (*6*)) to leverage an incredibly simple differential contextual fear conditioning paradigm (Fig. 1a).

**Figure 1.**
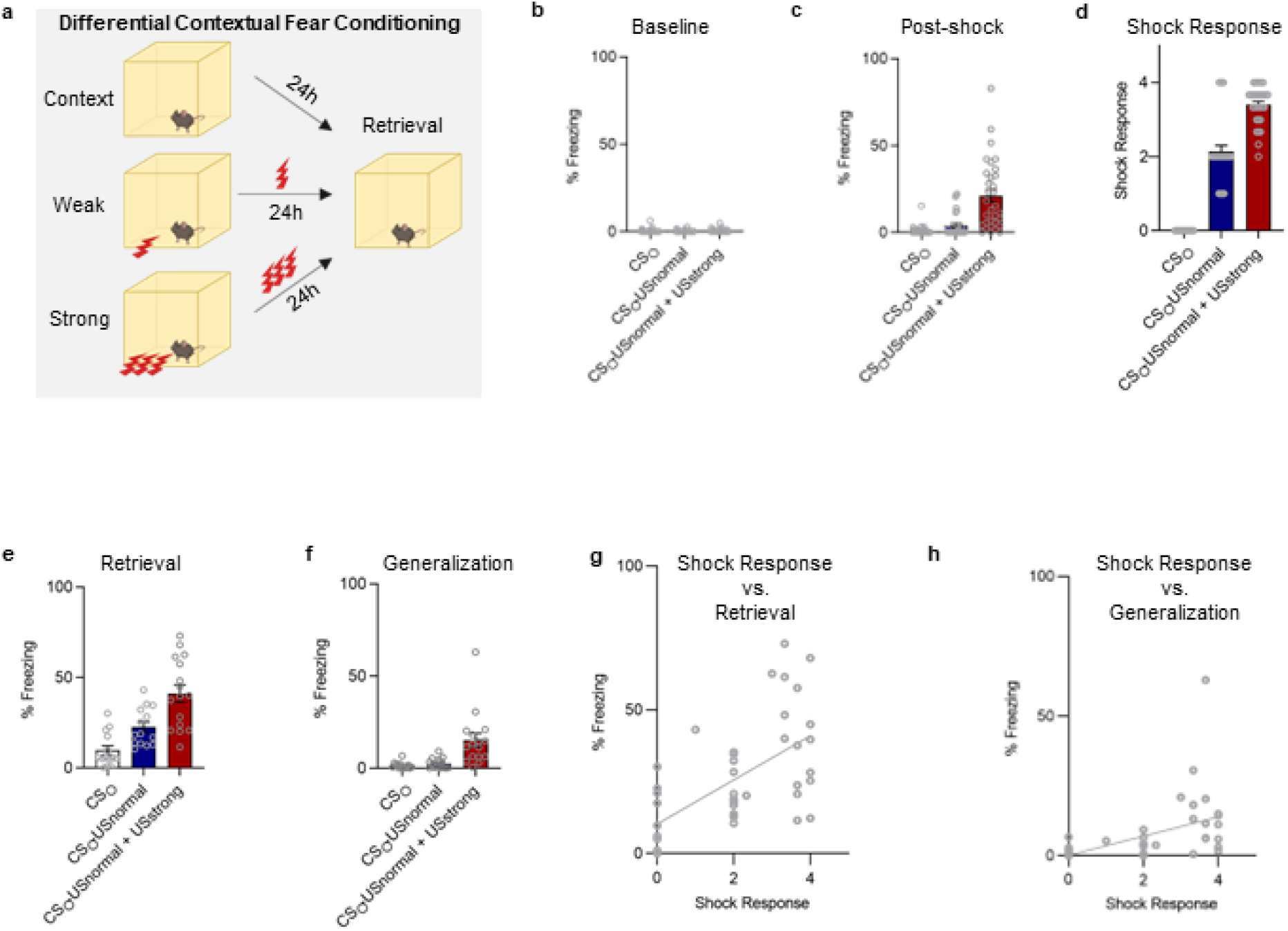
Stronger Conditioning produces stronger long-term memories that generalize to safe environments. (a) mice were trained in a differential fear conditioning paradigm where only the tactile US foot-shock. This differential contextual conditioning paradigm only differed only by the presence and intensity of foot shocks across three groups, CS_○_ (no foot shocks), CS_○_US_normal_ (1x 1s, 0.4mA foot shock) and CS_○_US_normal_ + US_strong_ (3x 1s, 1.0mA foot shocks), before being returned to the same context 24hr later to measure memory strength. (b) Mice showed no difference in baseline freezing, F_2,89_=0.5432, p=0.5828. (c) Mice significantly differed at post-shock freezing W_2,46.4_=16.63, p<0.0001. (d) Shock response scores were significantly different between all groups, H_3_=73.19, p<0.0001. Shock groups significantly differed from CS_○_ (p_adj._’s<0.0001), and CS_○_US_normal_ vs. CS_○_US_normal_ + US_strong_ differed (p_adj_=0.0045). (e) Groups significantly differed at a 24-hr retrieval test, W_2,26.67_=17.83, p<0.0001. (f) Groups significantly differed when tested in a new environment during generalization W_2,23.76_=7.055, p<0.0039. (g) Shock response scores were significantly correlated with % freezing at retrieval r(44)=0.6222, p<0.0001. (h) Similarly, shock response scores were significantly correlated with % freezing during generalization r(44)=0.4796, p=0.0010.

In contextual fear conditioning, the overall “contextual/configural” representation of an episodic memory is a product of the animal’s senses. Thus, real-world sensory inputs coming from auditory, visual, olfactory, gustatory, and incidental tactile sensations (i.e., components of the multisensory representation) can be treated as static given they have a quasi-stochastic sampling pattern across individuals; henceforth mathematically represented with subscript “○“). For example, suppose individual animals sample a neutral multisensory context in a quasi-random or stochastic nature (NS_○_). NS_○_ can be decomposed into individual sampling rates through a particular sensory modality (e.g., incidental tactile, visual, gustatory, olfactory, auditory) for a given moment in time (t). Therefore, we did not focus in on all of the potential permutations of a complex multisensory context-NS_○_ (e.g., *NS*^x1p(a)+x2p(v)+x3p(t)+x4p(o)+x5p(g)/t^, where the given state/value of NS_○_ is a sum of the weighted (x1, x2…x5) probability (p) of the magnitude (0 ≥ x ≤ 100%) of a given sensory input (auditory (a), visual (v), tactile (t) …) in a given moment of time “t” and represented by a distributed network of cells and circuits (*78*). We did focus on the single sensory input we can precisely control – the foot-shock unconditioned stimulus (US). Importantly, differences in the magnitude of the foot-shock US produces a reliable and replicable increase in the magnitude of unconditioned defensive response (UR). Using this logic, we could modify the foot-shock US on three domains (duration (d), intensity (i), and number (n)) in order to determine how different tactile foot-shock US inputs (over stochastic “○“ incidental tactile sensory inputs incorporated into the multisensory context-NS_○_ during memory acquisition) influences the strength of an episodic NS_○_+US→CS memory (e.g., a “fear” or threat memory).

Importantly, our recent work revealed that many USs (e.g., tactile, olfactory, or auditory) create neural activity patterns in the CA1 subfield of the ventral hippocampus that parse diverse unconditioned sensory inputs in a complex way (*79*). Despite the limitations of that study, we also found that temporal variations in a US, irrespective of the particular sensory modality engaged, creates elevated variability in calcium activity as measured by in vivo calcium recordings (*79*). These data suggested that the duration of a US must be held constant (*79*) in order for us to detect a direct biological component of the US during the formation of an episodic memory. Thus, the US which is incorporated into the multisensory contextual-NS_○_ (e.g., the NS_○_ + US association) can be represented by the formula: 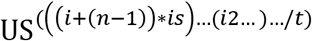, where “i” is intensity of a particular sensory input, (n-1) is the number of US presentations through a given sensory input, “i_s_” is the internal state of the animal (as yet to be defined), and “i2…i3…” are other USs from other senses constrained in the same way as “i” in the same given moment of time “t”. Thus, the discrete multisensory context-NS_○_ memory can be represented by the equation: 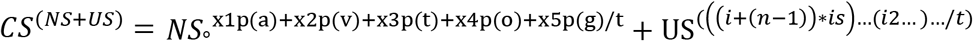, given the CS can be decomposed into a fundamental association between the stochastic multisensory contextual-NS_○_ and the US. Importantly, in this model, information about the modality specific US is still present and subtractable from the future CS association.

Using the above logic, we contextually fear conditioned three groups of animals by varying only two domains of the tactile foot-shock US: foot-shock intensity in mA and the number of actual foot-shocks presented across time thereby simplifying the 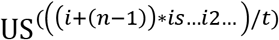 term to “weak” 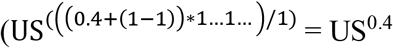 and “strong” 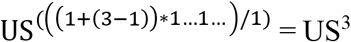, given mice were genetically similar and treated to have the same quasi-stochastic internal state during all aspects of the experience – behaviorally evident in baseline levels of freezing. Mice received either no shock and contextual exposure (Context; NS_○_ e.g., incidental learning control), a single mild foot-shock (NS_○_+US^weak^), or three strong foot-shocks (Strong; ((NS_○_+US^weak^) + US^strong^); Fig. 1a). Mice showed no difference in baseline levels of freezing (Fig. 1b), but mice receiving stronger conditioning (three 1-mA foot-shocks) exhibited higher post-shock freezing (e.g., a measure contentiously used as a proxy for acquisition) relative to other groups (Fig. 1c).

We also measured the active unconditioned response (UR) to the tactile foot-shock US inputs and detected a robust, reliable, and separable difference in the magnitude of the UR (Fig. 1d). Moreover, and as would be predicted, the additional intensity and number of USs at conditioning produced different strengths of an episodic memory – readout behaviorally as elevated defensive freezing to the conditioning context 24-h later (a conditioned stimulus or CS; Fig. 1e). As may also be predicted, additional USs at conditioning produced fear generalization to a novel environment 48-h later (Fig. 1f). Moreover, there was a significant correlation with US shock responses and defensive freezing at retrieval and generalization (Fig. 1g-h). Taken together with our recent work on the behavioral mechanisms of fear generalization (*8, 9*), these data suggest fear generalization to complex multisensory environments is likely a product of embedding additional US^strong^ information into the NS_○_+US^weak^ association in order to produce a NS_○_+US^weak^ + US^strong^ memory. Furthermore, these data show a strong relationship between the US input parameters (intensity x number), discrete UR motor output type (0-4 ordinal scale; no response flinch, hop, horizontal jump, vertical jump;(*80*)), and the later retrieval and expression of an episodic memory via a conditioned freezing response (CR) – a traditional proxy for the strength of a NS_○_+ US or “CS” memory.

Assuming 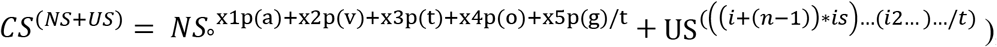, along with our recent work on the environmental constraints of recent and remote fear generalization (*8, 9*), we hypothesized that the added US^strong^ information must be separable from all other multisensory noise in the hippocampus at the biological level during the formation of a “strong” episodic memory given the UR / CR relationship above (*79*). By contrasting animals against the multisensory context-NS_○_ controls, we could reliably compare the biological bases of US^weak^ to that of US^weak^ + US^strong^ by subtracting out the stochastic multisensory context-NS_○_ sampling during conditioning. That is, NS_○_-US^weak^ - NS_○_ (context) = US^weak^ versus NS_○_+ US^weak^ + US^strong^ - NS_○_ = US^weak^ + US^strong^. We used this logic to examine how a novel class of RNA, Circular RNAs – which are remarkably stable and persist across time – may represent additional US^weak^ relative to US^weak^ + US^strong^ information, given US^strong^ was behaviorally evident 24-48h following conditioning (Fig. 1d-e). We focused in on the CA1 subfield of the dorsal hippocampus given our recent hippocampal work as well as the overwhelming evidence that the dorsal CA1 subfield in rodents is critical for storing episodic memories (*81*). Given normal variations in activity dependent RNA expression associated with memory, we collected brain tissue 1-hr after conditioning.

We hypothesized that circular RNAs may (1) operate in an activity-dependent manner and (2) parse the additive US^strong^ biological information embedded on top of the episodic NS_○_+US^weak^ (or CS) association during memory acquisition. We conditioned animals using identical parameters (Fig. 2a), collected brain tissue 1-hr after conditioning, and carefully dissected out bilateral punches of the CA1 subfield in the hippocampus – given this area is critical for memory and it is where the multisensory contextual NS_○_ representation is stored early after learning (Fig. 2b). We performed next-generation sequencing for circular RNAs on pooled samples across all cells of the dorsal hippocampus CA1 subfield given difficulty in detection without abundant RNA (Fig. S1–2). The need for significant input RNA also highlights the low activity-dependent abundance of circular RNAs in CA1 given brain tissue is not known to contain the key RNAses.

**Figure 2.**
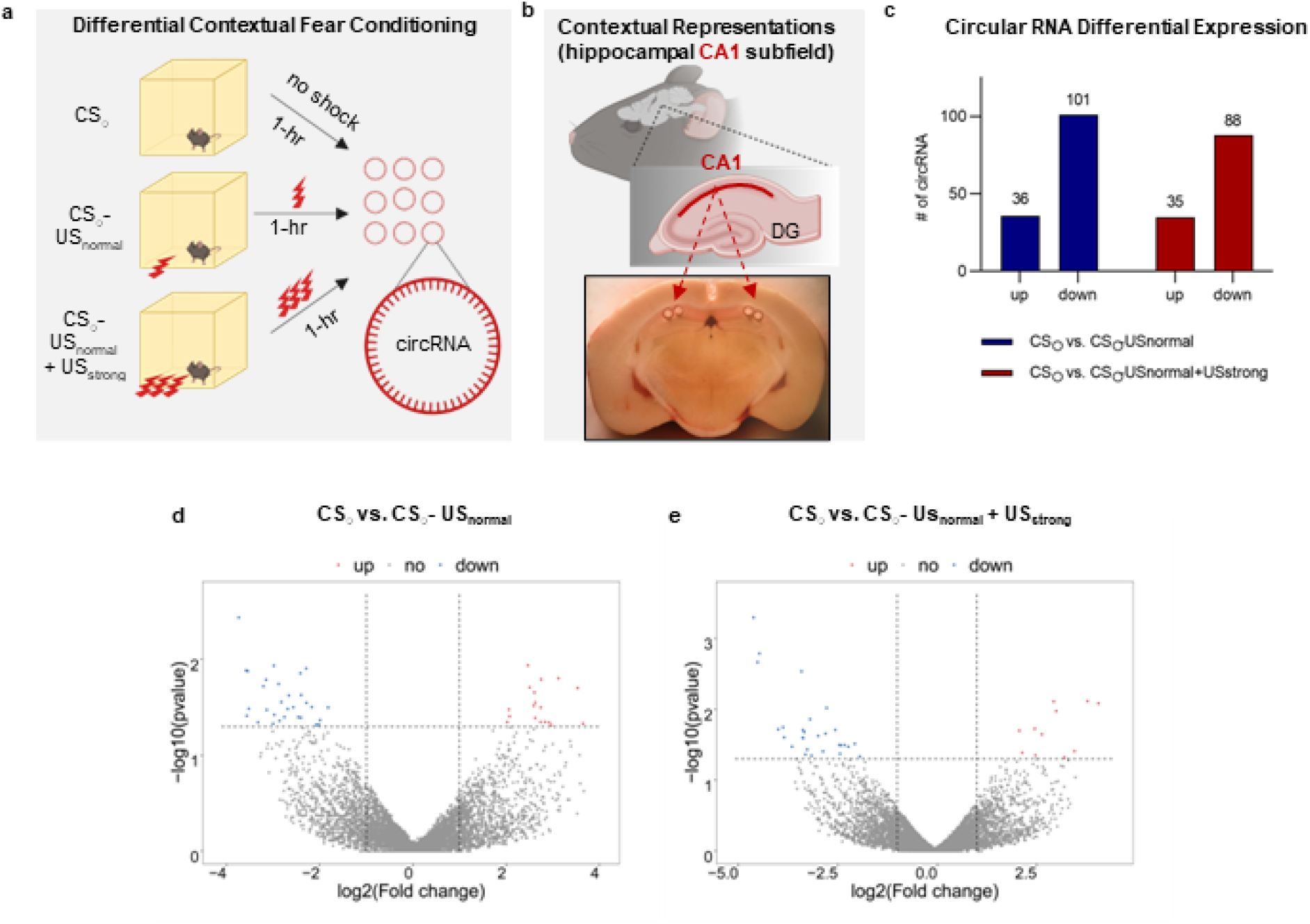
Circular RNAs are regulated in an activity-dependent manner in the CA1 subfield of the dorsal hippocampus. (a) Overview of behavioral paradigm with CS_○_ (context), CS_○_US_normal_ (weak), and CS_○_US_normal_+US_strong_ (strong). One after exposure to the context, weak training, or strong training, brains were removed. (b) brains were sectioned on a cryostat and two 0.5mm punches were taken from each hemisphere. (c) Groups were compared against the context-only exposed controls. CS_○_US_normal_ mice showed 36 up-regulated and 101 down-regulated Circular RNAs at 1-hr following conditioning. CS_○_US_normal_+US_strong_ mice showed 35 up-regulated and 88 down-regulated Circular RNAs at 1-hr following conditioning. (d-e) Volcano plots of up (blue) and down (red) Circular RNAs in CS_○_US_normal_ (left) or CS_○_US_normal_+US_strong_ (right).

Circular RNA sequencing 1-hr after conditioning revealed a remarkably small number of differentially expressed circular RNAs that were regulated in an activity-dependent US^weak^ vs. US^strong^ manner. Thus, US^weak^ mice showed an up-regulation of only 36 circRNAs and down-regulation of 101 circular RNAs (Fig. 2c). Surprisingly, US^strong^ mice showed an upregulation of only 35 circular RNAs and a down-regulation of 88 circular RNAs (Fig. 2c). Thus, only a single upregulated circular RNA differed between the US^weak^ and US^strong^ groups – which were identical (Fig. 2a) to every aspect of an episodic memory, except for the intensity and number of tactile USs. Given the low abundance, we factored in overall transcript number on top of statistical analysis to identify that the top up-regulated circRNA for US^weak^ was circRNA_13680 related to syntrophin, basic-2 (Sntb2) whereas the 2^nd^ most up-regulated hit was circRNA_1766 related to the RAB3 GTPase protein subunit 2 (RAB3). It should be noted we also detected the greatest downregulation of a circular RNA related to the RNA-binding protein Pumillo1 (Fig. S2–3; (Pum1; (*31*)).

For US^strong^, circRNA_13082 related to alkaline ceramidase-3 (Acer3) was the most abundant up-regulated target followed by circRNA_12350 related to mitogen activated protein kinase 14 (MapK14). By contrast, circRNA_2941, also related to MAPK14, was the most down-regulated target. Importantly, each of the genes represented by these particular Circular RNA variants (e.g., Sntb2, Rab3b, Pum1, Acer3, or MapK14) contain a large number of variants (see (*82*)) and have been implicated in different aspects of cellular plasticity or learning and memory.

Surprisingly, US^strong^ (Fig. 3a) exhibited an elevation in Circular RNA gene ontologies associated with protein kinase activity and protein phosphorylation, as well as tyrosine kinase signaling – key biological processes of long-term memory (Fig. 3b; (*38, 83–85*)). By contrast, the US^weak^ conditioned group showed an elevation in Circular RNA gene ontologies associated with protein de-phosphorylation and intracellular signaling – key biological processes associated with weaker, shorter-lasting forms of memory (Fig. 3c; (*5, 39*)).

**Figure 3.**
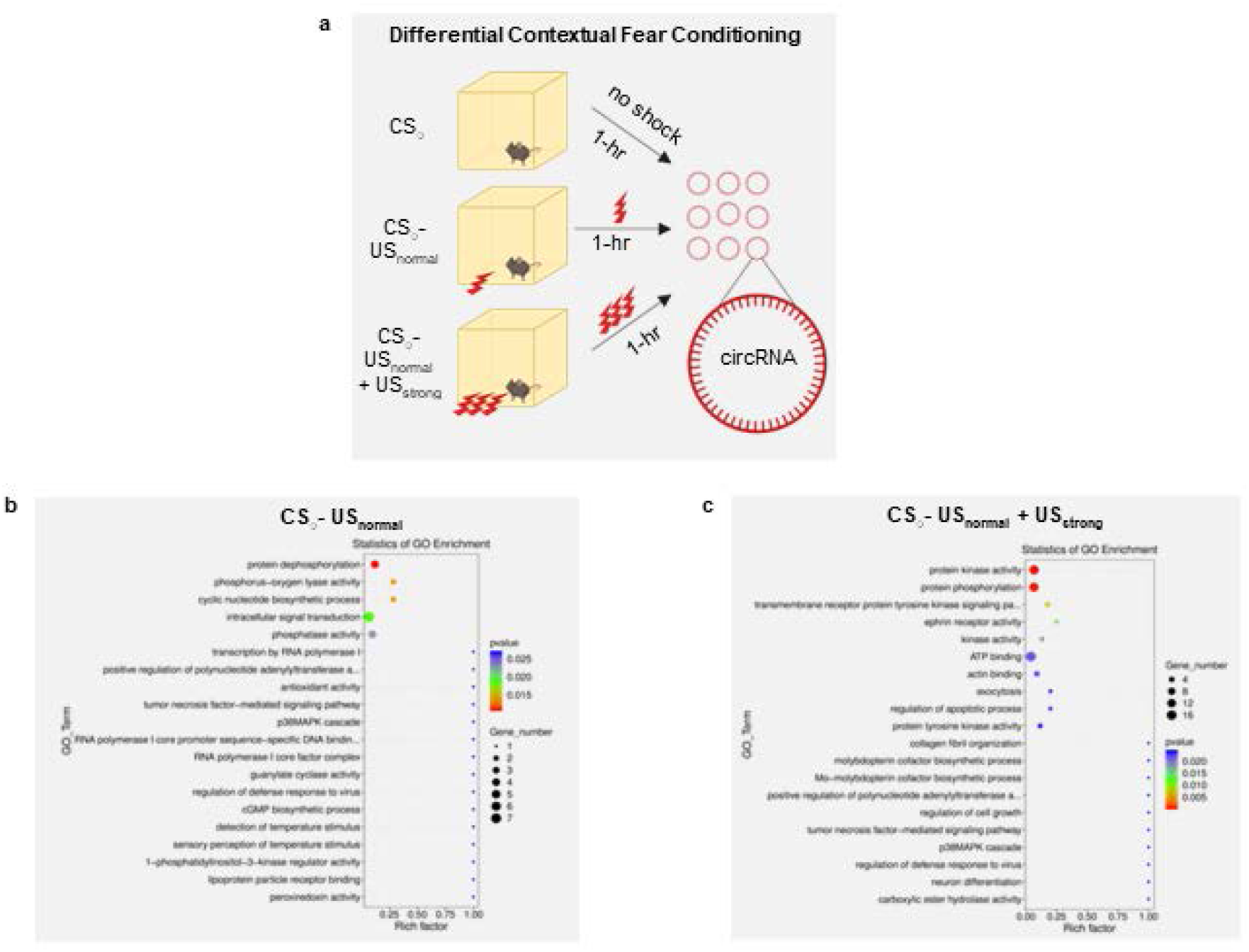
Gene ontologies of Circular RNAs. (a) schematic of behavioral paradigm. (b) context-only exposed versus weakly exposed animals. (c) context-only exposed animals versus strongly conditioned animals. Individual points are color coded by the p-value and represent the enrichment value. Enrichment value represents normalization (z) and was calculated by: z_sample_ = (log_2(individual sample)_ – log_2 (mean of all samples)_ / log_2 (standard deviation of all samples)_.

Taken together, these data show a remarkable parallel between the identified Circular RNAs and well-known gene/gene families related to long-term memory (*5*). Thus, given their known biological stability, we investigated the presence of Snt2b in US^weak^ to visually confirm the presence and density of Circular RNAs in the CA1 subfield of the hippocampus. Snt2b was densely clustered in the CA1 subfield (Fig. S5). Given the density, these data suggest that Circular RNAs may be present in cells of a memory trace.

Considering these data, we propose a new framework for multisensory configural representations whereby forward-spliced RNA (e.g., mRNA, ncRNA, etc.) and proteins act to strengthen synaptic connections, circuits, and networks for a particular multisensory CS_○_ memory and backward-spliced Circular RNAs act to represent a discrete US within that multisensory configural representation (Fig. 4). Thus, this dual-process biological model of multisensory representations or episodic memory provides a framework for better deciphering the components of complex associations formed in the real-world.

**Figure 4.**
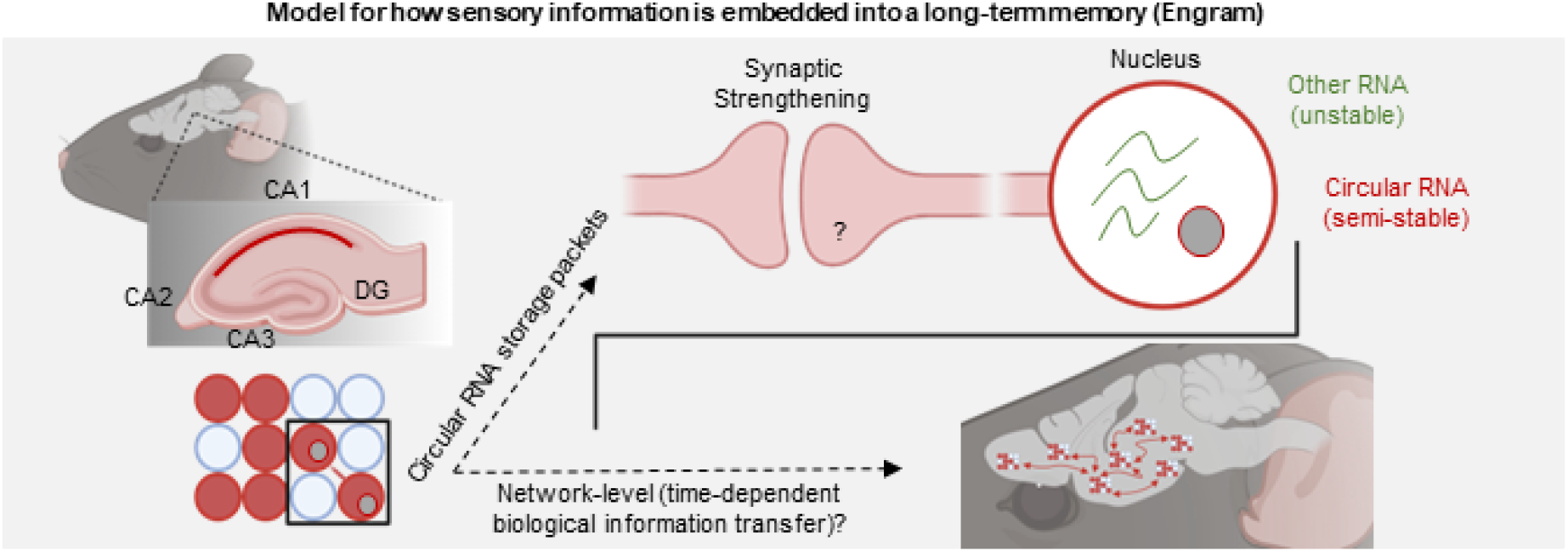
Revised model for long-term memory. Circular RNAs provide a stable storage mechanism for real-world sensory US information. Several questions remain denoted with a question mark. (1) What role do Circular RNAs have in modulating the strength of CS-US associations at the synapse. (2) Do Circular RNAs have a time-dependent role in systems consolidation for a particular memory?

## Discussion

Our data show that a novel class of RNA, Circular RNAs, are associated with the formation and storage of a long-term episodic memory in the brain. First, we expand on simple Pavlovian frameworks of the 20^th^ century to mathematically constrain how complex multisensory environments can be associated with unconditioned stimuli in the real-world using the formula: 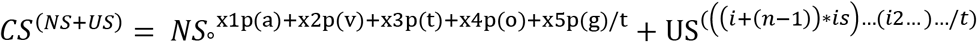. Second, we leveraged this framework to identify how fear generalization under binary conditions (e.g., US^weak^ < 1 and US^strong^ > 1) is associated with embedding additional US information. Third, we used next-generation sequencing to identify how a novel, persisting class of RNA, Circular RNAs, are enriched in an “activity-dependent” US^weak^ versus US^strong^ manner 1-hr after conditioning in the CA1 subfield of the dorsal hippocampus (CA1). A timepoint beyond the traditional transcriptional wave associated with learning and memory. Fourth, we used in situ hybridization to visually confirm the presence and density of Snt2b in CA1.

Pavlovian accounts of learning that emerged in the early 20^th^ century established rudimentary mathematical principles of associative learning. Indeed, in the latter half of the 19^th^ century, our laboratory leveraged these principles along with a radically reductionist approach to decipher how long-term memories involve synaptic strengthening and discrete cellular/molecular cascades (cf. (*5, 6*). Studies dating back to the 1970s have found higher US intensities drive greater long-term defensive responses (e.g., startle, freezing), fear generalization, and “dose”-dependently drive increased transmitter levels in the brain across species (*86–89*). Several other theories have also addressed aspects of associability (*67*). These studies suggest that the US plays an especially important role in regulating how a neutral stimulus (NS) is associated with an unconditioned aversive stimulus (US) to produce a conditioned stimulus (CS) and conditioned defensive response (CR). More research is needed to understand how USs, URs, and CRs are behaviorally related in complex environments in addition to how USs are biologically represented in distinct and overlapping memory traces (*90*).

Our refined mathematical framework synthesizes key principles of associative learning and the real-world to provide a simple account for how complex multisensory episodic memory traces/states are biologically embedded in the brain. Although our model accounts specially for fear generalization, it does not account more generally for valence (*91*). Thus, future work should examine how valence is biologically encoded within the US in addition to how discrete parametrizations of different internal states (is) influence how the US is biologically incorporated into memories in the weeks and months following the initial storage.

By leveraging this revised mathematical framework along with two different methods (i.e., next-generation sequencing and in situ hybridization), we show the presence of key Circular RNAs 1-hr after conditioning which may persist longer. Our data suggest that different sets of Circular RNAs are regulated in (1) an activity-dependent and (2) intensity-specific manner.

It has long been appreciated that the hippocampus is essential for memory (*58, 92*). Our findings highlight a new piece to the long-standing puzzle of memory whereby we may finally close the loop. That is, forward-spliced synaptic plasticity in neuronal ensembles active during the initial time of learning, represent a particular multisensory configural state (*7, 18, 93*). By contrast, backward spliced Circular RNAs likely contain lasting biological information related to real-world USs. More research is needed to identify how Circular RNAs representing different USs can be transferred between organisms as well as how epigenetic modifications and phase-separated states modulate information storage over time.

One of the biggest critiques to the notion of RNA as a biological storage mechanism for long-term memory is that different species of RNA (e.g., piRNAs, miRNAs, mRNAs, ncRNAs, etc.) are unstable and easily degraded. Thus, traditional RNAs do not persist on time scales relevant to episodic memory storage. Moreover, these challenges in stability are most evident in the current class of next-generation mRNA-based therapeutics that emerged during the COVID pandemic that are encapsulated inside of nanoparticles to enhance stability and delivery (*94*). Circular RNAs overcome these limitations. Although we show that Circular RNAs are present 1-hr after learning, it is likely they are stable for much longer time scales. More research is needed to identify how different USs (which are generally conserved across mammalian species) are encoded by different Circular RNAs, as well as the timescales on which Circular RNAs persist following learning.

Research in the coming decades should focus on the molecular biology of Circular RNAs in relation to how USs are biologically incorporated into associative memories to determine their strength, as well as their role in transgenerational inheritance of sensory information and several other forms of memory. Moreover, the presence and possible long-term stability of Circular RNAs may provide a new means for targeting the strength or content of multisensory representations or episodic memories. Indeed, Circular RNAs may provide a novel target and treatment strategy across several pathological conditions such as Alzheimer’s disease, post-traumatic stress disorder, anxiety, and many more (*60*).

## Acknowledgements

I thank Eric Kandel for his initial thoughts on the manuscript. I thank Christina Doyle for help in acquiring reagents and overall laboratory support. I thank David Glanzman for casual discussions at the 2019 Society for Neuroscience meeting.

## Author contributions

A.A. conceived of the study. A.A. performed the behavioral procedures. A.A. refined protocols. A.A. wrote the manuscript.

## Methods

### Animals

10–18-week-old C57/BL6 mice were used for all experiments (Jackson Laboratory, Bar Harbor, ME). Same sex male mice were housed 4 to a cage on a 12h light/dark cycle (lights on from 7:00 – 19:00) with ad libitum access to food and water. Mice were acclimated to the colony for at least 1-week prior to the start of experimentation.

All experimental sessions were conducted during the light phase between 09:00 and 17:00 h. Procedures were conducted in accordance with the US National Institute of Health Guide for the Care and Use of Experimental Animals and were approved by the Columbia University Institute of Comparative Medicine at the Zuckerman Mind Brain & Behavior Institute at Columbia.

### Behavior – Tail & Transport Habituation (Days 1-3)

On day 1, animals were tail-marked (not ear-tagged) with a non-toxic marker to expose animals to tactile stimulation of the tail and uniquely identify each animal. On days 2-3, animals were transported to the fear conditioning room, positioned on a holding rack, and food removed. Mice were habituated to the conditioning room on the holding rack for 1-2 hrs/day.

### Behavior – Fear Conditioning (Day 4)

On day 4, we used identical transportation procedures to days 2-3. After 1-hr of habituation, mice were conditioned using a standard 4-chamber NIR video fear conditioning system (Med Associates Inc., St. Albans, VT). Individual conditioning chambers measured. Chambers were identical spatial (standard square layout), olfactory (70% ethanol), visual (white lighting), tactile (metal grid-floors) during conditioning. Freezing was measured via integrated Med Associates Video FreezeView software with an episode of freezing defined as immobility for 75% of 30 frames in a 1-s time bin. For each training session, 1 cage (4 animals) was transported next to the conditioning chambers. Mice were lifted by their tail and placed in the chamber. Following placement of the last animal, the session was started.

For context-only incidental learning controls (“context”), mice were exposed to a 120-s baseline period (“baseline”) followed by non-reinforced exposure to the camber for the same time as “weak” or “strong” conditioning (180-s or 210-s). “Context” mice were collapsed together across all experiments.

For “weak” conditioning, animals were exposed to a 120-s baseline period (“baseline”) followed by a single 1-s 0.4-mA foot-shock, followed by a 60-s non-reinforced period (“post-shock / acquisition”).

For “strong” conditioning, animals were exposed to a 120-s baseline period (“baseline”) followed by 3 1-s 1-mA foot-shocks spaced 10-s apart, followed by a 60-s non-reinforced period (“post-shock / acquisition”).

Following behavior, each cage was returned to a new holding rack. Animals were left on the holding for 30-m following behavior with the last cage of mice and then returned to the same rack locations in the colony.

### Behavior – Fear Retrieval (Day 5)

On day 5, mice were returned to the conditioning room and habituated like days 2-3. All groups, “context”, “weak”, and “strong” were exposed to the original training boxes and chambers in a counter-balanced test order relative to conditioning. Chamber were cleaned with 70% ethanol.

### Behavior – Fear Generalization (Day 6)

On day 5, mice were returned to the conditioning room and habituated like days 2-3. All groups, “context”, “weak”, and “strong” were exposed to different test boxes and chambers that differed along, spatial (circular chamber walls), olfactory (1% acetic acid), visual (NIR lighting), tactile (opaque acrylic floor) components. Mice were tested in a counter-balanced test order relative to conditioning. Following behavior, each cage was returned to a new holding rack. Animals were left on the holding for 30-m following behavior with the last cage of mice and then returned to the same rack locations in the colony.

### Tissue Collection (1-hr and 4-days following conditioning)

Brains were collected using methods identical to our previous work (*77*). Briefly, 1-hr or 4-days following conditioning, mice were transported to a room adjacent to the colony. Brains were rapidly removed, and flash frozen in isopentane on dry ice and stored at −80°C until sectioning.

Brains were sliced on a cryostat at −24°C. An area corresponding to the dorsal hippocampus which represented all subfields including area CA1 was targeted (−1.94 Bregma). Upon reaching these coordinates, 0.35mm (diameter) x 0.5mm (depth) tissue punches of area CA1 OR 16-20um tissue sections were collected. For tissue punches, two adjacent tissue punches targeting area CA1 were collected bilaterally (4 punches total) and immediately placed into a −80°C 1.5mL microcentrifuge tube for circular RNA sequencing. For tissue sections, 4 serial sections were collected onto a single charged microscope slide and stored in a slide box at −8°0C prior to in situ hybridization.

### Circular RNA sequencing

Total RNA was extracted with Trizol (Invitrogen, CA, USA). Total RNA quantity and purity were analyzed of Bioanalyzer 2100 and RNA 6000 Nano LabChip Kit (Agilent, CA, USA) with RIN > 7.0. Approximately 10 ug of total RNA was used to remove ribosomal RNA using an Epicentre Ribo-Zero Gold Kit (Illumina, San Diego, USA). Linear RNA was digested following standard protocols using Epicentre Ribonuclease R (Illumina, San Diego, USA). Following purification, the linear RNA fractions were fragmented into small pieces with divalent cations under heat. The cleaved RNA fragments were reverse-transcribed to create the final cDNA library in accordance with a strand-specific library preparation using the dUTP method. The average insert size for the paired-end libraries was 300 ± 50 bp. Sequencing was conducted using a paired-end 2× 150bp sequencing reaction on an Illumina Hiseq 4000 sequencer.

The following bioinformatic pipeline was used to identify circRNAs: (1) Quality control, FastQC (v.0.10.1); (2) Adapter trimming, Cutadapt (v.1.10); (3) Genome alignment, Tophat (v.2.0.4); (4) Back splice junction filtering, Tophat-fusion (v.2.1); (5) CircRNA identification, CIRCExplorer (v. 2.2.6) and CIRI (v.2.0.2); (6) Differential expression analysis, edgeR (v.N/A).

### Circular RNA in situ Hybridization

Tissue sections were processed for RNAscope in situ hybridization following removal from the −80°C slides were immediately fixed with 10% NBF for 15-min at 4C. Next, slides were rinsed in 1X PBS+DEPC at RT followed by 0.25% acetic anhydride and 0.1M TEA (pH 8.0) for 10 min. Slides were then successively washed in 50%, 70%, and 100% ethanol for 5-min then air dried for 5-min. Tissue sections were then treated with RNAscope hydrogen peroxide at RT for 10-min followed by rinses in 1x PBS and then incubated with Pretreat 4 for 25-min at RT. Slides were then rinsed in 1x PBS prior to hybridization.

Sections were incubated with Probes targeting Acer 3, Snt2b, as well as standard positive and negative RNAscope control slides at 40C for 2-hrs and following standard protocols outlined by ACDbio.

### Imaging

Images were captured on an Olympus FV1000 confocal microscope or a Keyence BZ-X800. Schematics in figures were created using Biorender and Microsoft Powerpoint.

### Statistics

Homogeneity of variance was first tested between groups and tests were corrected for any violations. For behavior 2 outliers greater than 2 standard deviations were removed from analyses. For behavioral analyses, Bartlett’s test was first used to test for homogeneity of variance. Welch’s test was used for comparison followed by Tukey’s post-hoc tests. For shock responsivity testing, Kruskal-Wallis tests were used followed by Dunn’s multiple comparisons. For RNA sequencing, standard Edge-R differential expression analysis was conducted using standard methods.

---

**Supplementary Figure 1.**
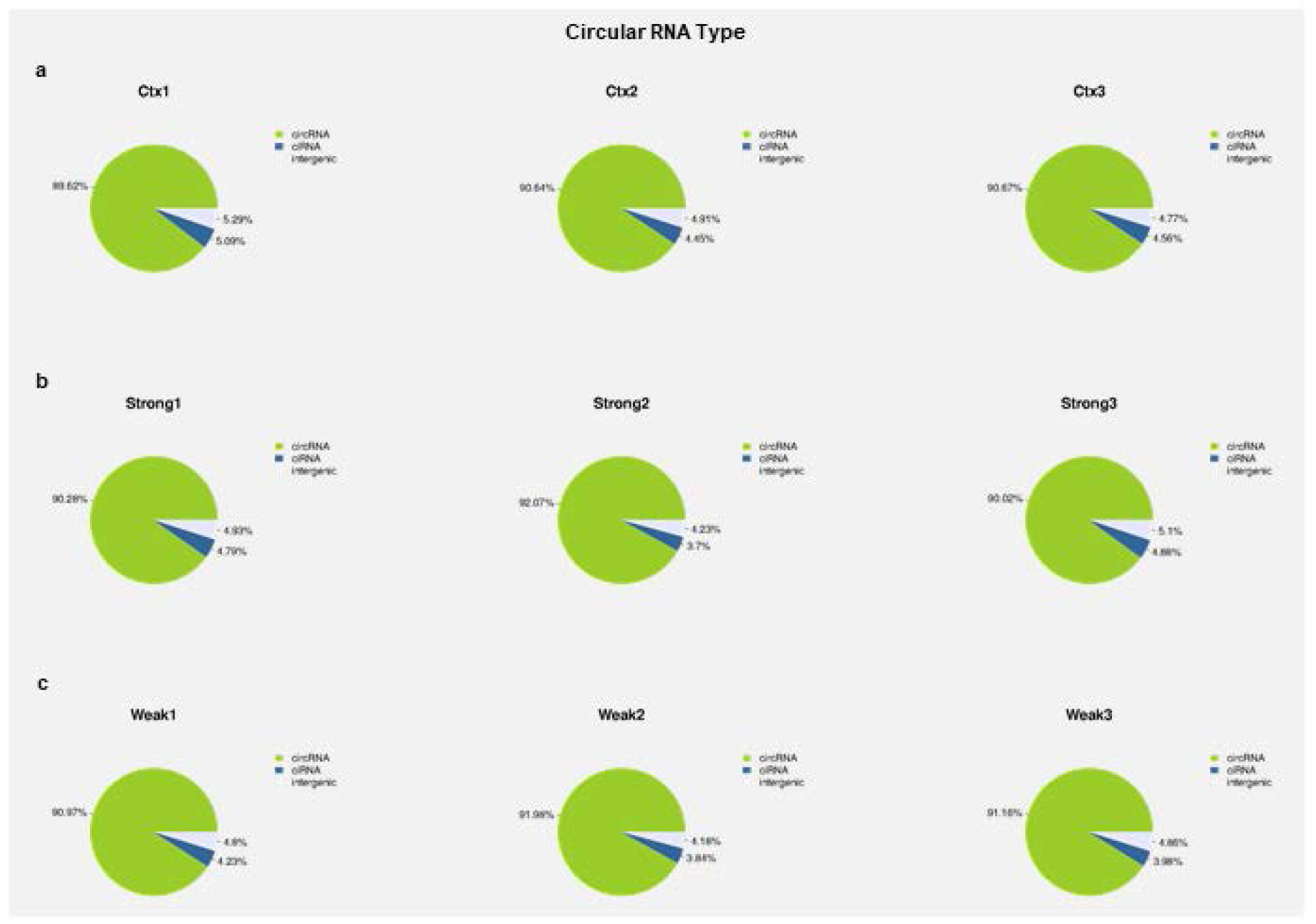
Statistics of different Circular RNA types across samples. Circular RNAs were mapped across different genomic segments to identify the proportion associated with exonic regions (green), intronic regions (blue), or intergenic regions (grey). (a) Context-only exposed mice statistics. (b) Strongly conditioned mice statistics. (c) Weakly conditioned mice.

**Supplementary Figure 2.**
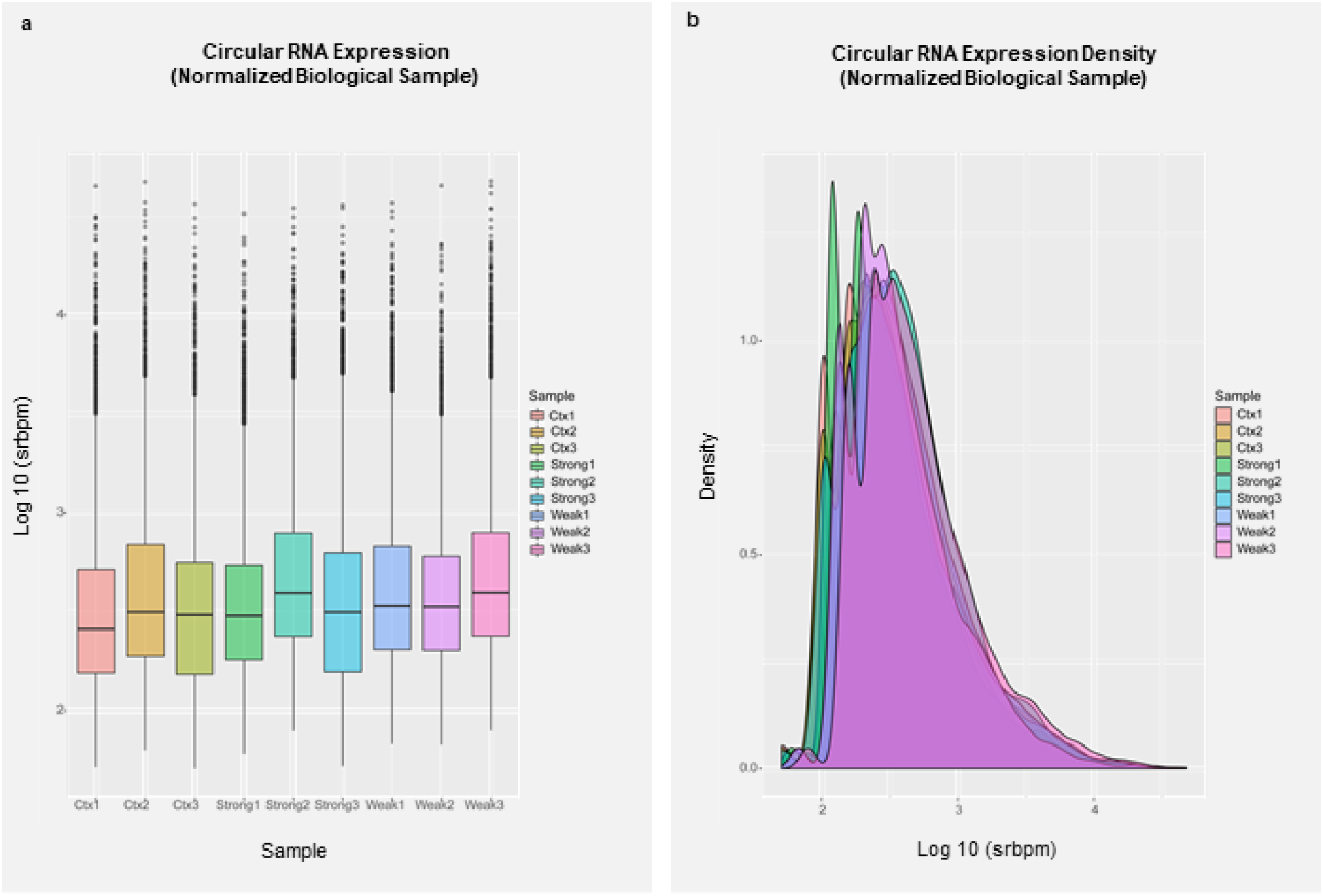
Normalized distribution of Circular RNAs. (a) boxplots of normalized Circular RNA expression in each biological replicate against spliced reads per billion mapped (srpbm). (b) Normalized Circular RNA expression density in each biological replicate.

**Supplementary Figure 3.**
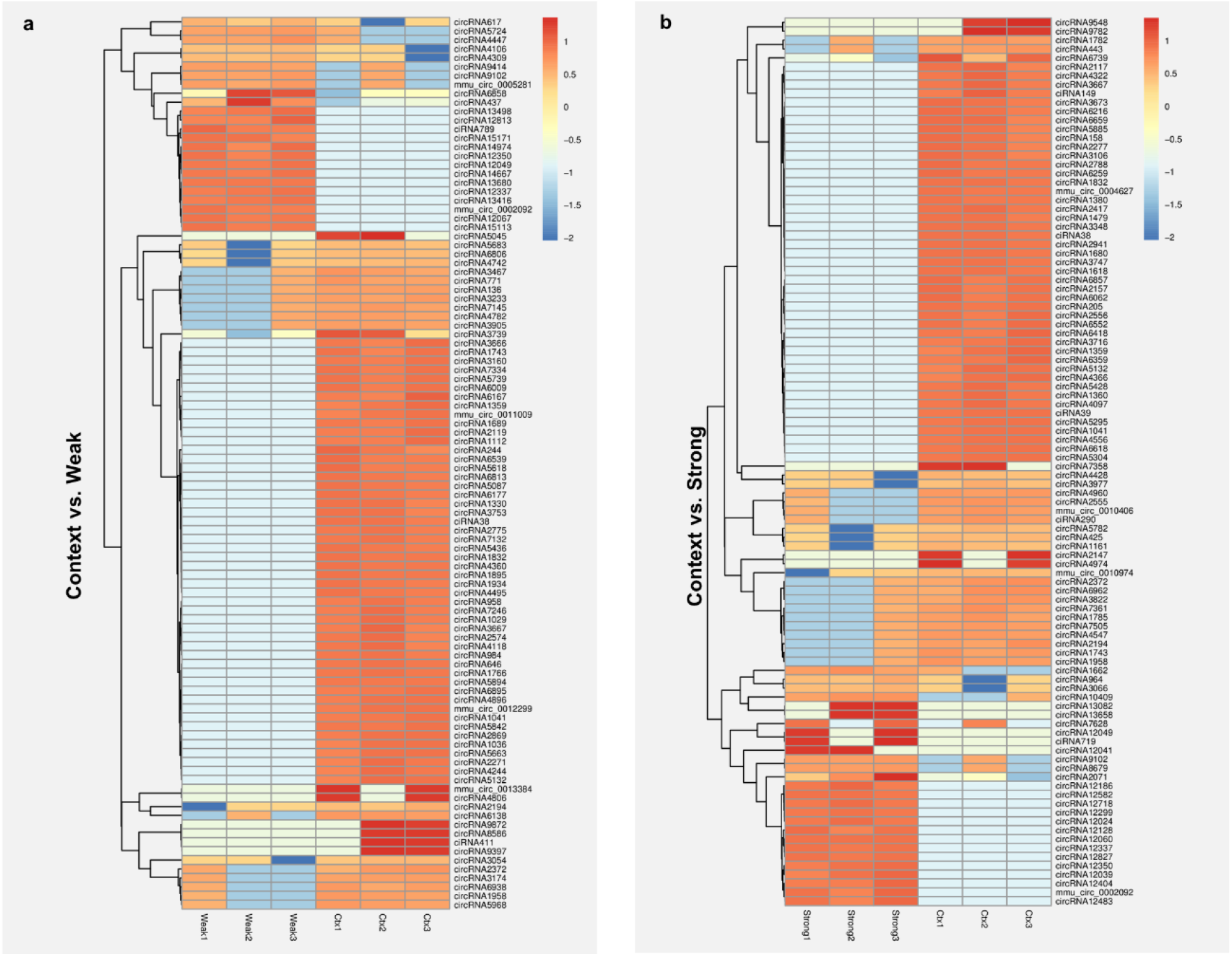
Heatmap of normalized differentially expressed Circular RNA reads in fragments per kilobase of exon per million mapped fragments (FKPM). (a) Context-exposed versus weakly trained animals. (b) context-exposed versus strongly trained animals. Normalization (z) was calculated by: z_sample_ = (log_2(individual sample)_ – log_2 (mean of all samples)_ / log_2 (standard deviation of all samples)_.

**Supplementary Figure 4.**
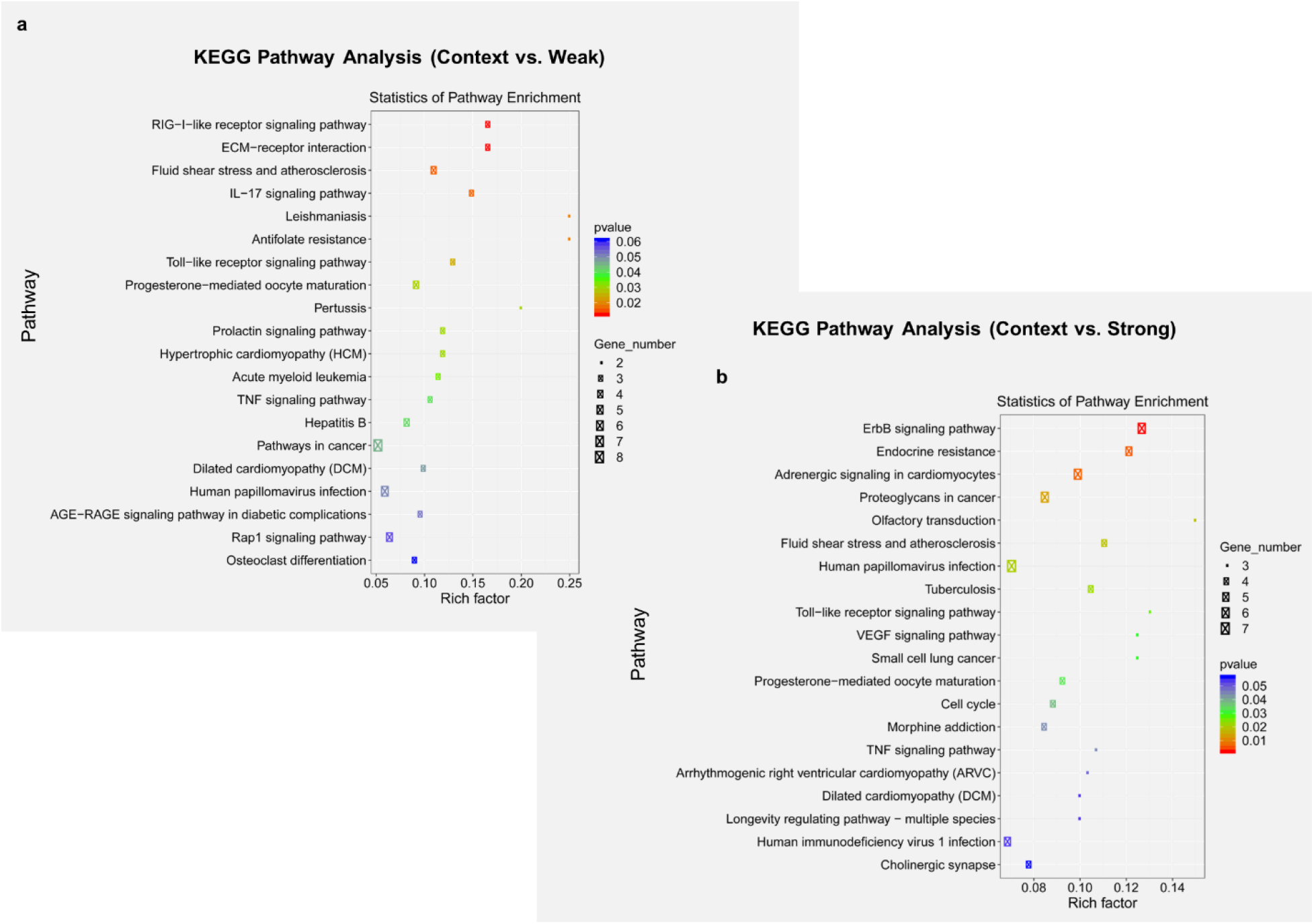
KEGG Pathway Analysis. (a) Context-exposed relative to weakly trained mice. (b) Context-exposed relative to strongly trained mice. P-values are color coded. Rich (enrichment) Factor = # of differentially expressed genes in the KEGG pathway / Total number of genes in KEGG pathway.

**Supplementary Figure 5.**
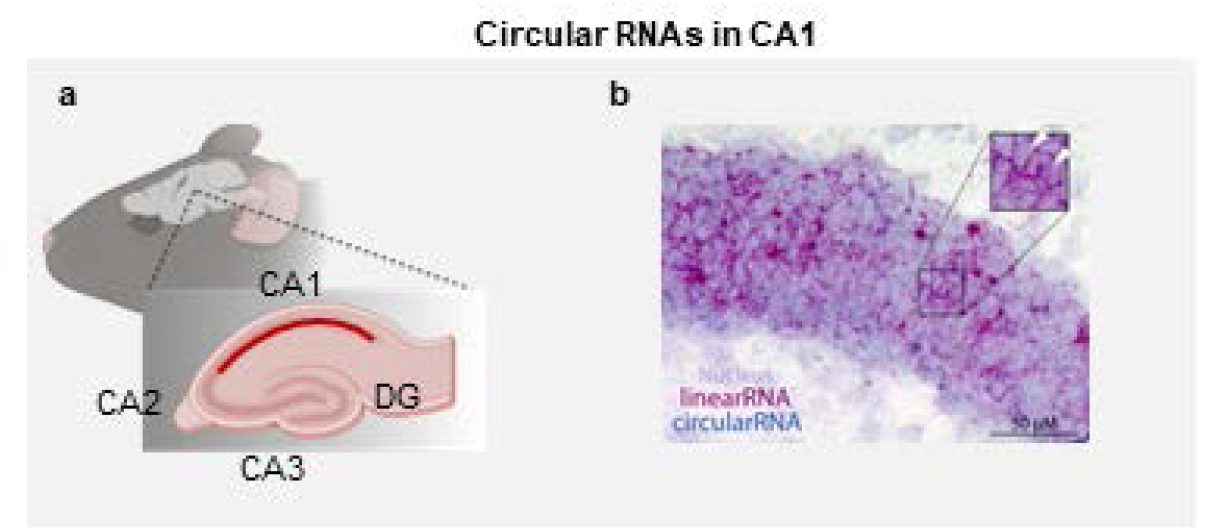
Preliminary in situ hybridization staining. (a) schematic of hippocampal subdivisions. (b) in situ hybridization for Snt2b in CA1. The nucleus is represented in purple, linear RNA depicted in red, and Circular RNA in blue.

## References

1. E. R. Kandel, In search of memory: The emergence of a new science of mind. (WW Norton & Company, 2007).

2. Plato, Plato’s The Republic. (Books, Inc., New York, 1943).

3. J. Beare, On memory and reminiscence Aristotle (ca. 350 bc). Annals of Neurosciences 17, 87 (2010).

4. M. Mayford, S. A. Siegelbaum, E. R. Kandel, Synapses and memory storage. Cold Spring Harbor Perspectives in Biology 4, (2012).

5. E. R. Kandel, The molecular biology of memory storage: a dialogue between genes and synapses. Science 294, 1030–1038 (2001).

6. A. Asok, F. Leroy, J. B. Rayman, E. R. Kandel, Molecular mechanisms of the memory trace. Trends in neurosciences 42, 14–22 (2019).

7. S. Tonegawa, X. Liu, S. Ramirez, R. Redondo, Memory engram cells have come of age. Neuron 87, 918–931 (2015).

8. A. Asok, E. R. Kandel, J. B. Rayman, The neurobiology of fear generalization. Frontiers in behavioral neuroscience 12, 329 (2019).

9. A. Asok et al., Sex differences in remote contextual fear generalization in mice. Frontiers in behavioral neuroscience 13, 56 (2019).

10. D. Mobbs et al., Viewpoints: Approaches to defining and investigating fear. Nature neuroscience 22, 1205–1216 (2019).

11. R. Adolphs, The biology of fear. Current biology 23, R79–R93 (2013).

12. D. J. Anderson, R. Adolphs, A framework for studying emotions across species. Cell 157, 187–200 (2014).

13. J. LeDoux, Rethinking the emotional brain. Neuron 73, 653–676 (2012).

14. J. Panksepp, Affective neuroscience: The foundations of human and animal emotions. (Oxford university press, 2004).

15. K. Abdou et al., Synapse-specific representation of the identity of overlapping memory engrams. Science 360, 1227–1231 (2018).

16. L. G. Reijmers, B. L. Perkins, N. Matsuo, M. Mayford, Localization of a stable neural correlate of associative memory. Science 317, 1230–1233 (2007).

17. X. Liu et al., Optogenetic stimulation of a hippocampal engram activates fear memory recall. Nature 484, 381–385 (2012).

18. S. Ramirez et al., Creating a false memory in the hippocampus. Science 341, 387–391 (2013).

19. A. R. Garner et al., Generation of a synthetic memory trace. Science 335, 1513–1516 (2012).

20. C. A. Denny et al., Hippocampal memory traces are differentially modulated by experience, time, and adult neurogenesis. Neuron 83, 189–201 (2014).

21. K. Z. Tanaka et al., Cortical representations are reinstated by the hippocampus during memory retrieval. Neuron 84, 347–354 (2014).

22. R. C. O’Reilly, J. W. Rudy, Conjunctive representations in learning and memory: principles of cortical and hippocampal function. Psychological review 108, 311 (2001).

23. F. Crick, Neurobiology: Memory and molecular turnover. Nature 312, 101 (1984).

24. J. Lisman, The CaM kinase II hypothesis for the storage of synaptic memory. Trends in neurosciences 17, 406–412 (1994).

25. T. Abel, K. C. Martin, D. Bartsch, E. R. Kandel, Memory suppressor genes: inhibitory constraints on the storage of long-term memory. Science 279, 338–341 (1998).

26. T. Abel et al., Genetic demonstration of a role for PKA in the late phase of LTP and in hippocampus-based long-term memory. Cell 88, 615–626 (1997).

27. E. D. Roberson, J. D. Sweatt, A biochemical blueprint for long-term memory. Learning & Memory 6, 381–388 (1999).

28. D. S. Ling et al., Protein kinase Mζ is necessary and sufficient for LTP maintenance. Nature neuroscience 5, 295–296 (2002).

29. T. C. Sacktor, How does PKMζ maintain long-term memory? Nature Reviews Neuroscience 12, 9–15 (2011).

30. K. Si, S. Lindquist, E. R. Kandel, A neuronal isoform of the aplysia CPEB has prion-like properties. Cell 115, 879–891 (2003).

31. J. B. Rayman, E. R. Kandel, Functional prions in the brain. Cold Spring Harbor perspectives in biology, (2016).

32. L. Fioriti et al., The persistence of hippocampal-based memory requires protein synthesis mediated by the prion-like protein CPEB3. Neuron 86, 1433–1448 (2015).

33. K. Si, E. R. Kandel, The role of functional prion-like proteins in the persistence of memory. Cold Spring Harbor perspectives in biology 8, a021774 (2016).

34. J. B. Rayman et al., Genetic perturbation of TIA1 reveals a physiological role in fear memory. Cell reports 26, 2970–2983. e2974 (2019).

35. R. Rahman, W. Xu, H. Jin, M. Rosbash, Identification of RNA-binding protein targets with HyperTRIBE. Nature protocols 13, 1829–1849 (2018).

36. E. D. Pastuzyn et al., The neuronal gene Arc encodes a repurposed retrotransposon Gag protein that mediates intercellular RNA transfer. Cell 172, 275–288 (2018).

37. J. Ashley et al., Retrovirus-like Gag protein Arc1 binds RNA and traffics across synaptic boutons. Cell 172, 262–274 (2018).

38. I. Jin et al., Anterograde and retrograde signaling by an Aplysia neurotrophin forms a transsynaptic functional unit. Proceedings of the National Academy of Sciences 115, E10951–E10960 (2018).

39. H. Bito, K. Deisseroth, R. W. Tsien, CREB phosphorylation and dephosphorylation: a Ca2+-and stimulus duration–dependent switch for hippocampal gene expression. Cell 87, 1203–1214 (1996).

40. R. J. Kelleher III, A. Govindarajan, H.-Y. Jung, H. Kang, S. Tonegawa, Translational control by MAPK signaling in long-term synaptic plasticity and memory. Cell 116, 467–479 (2004).

41. I. M. Nooren, J. M. Thornton, Diversity of protein–protein interactions. The EMBO journal 22, 3486–3492 (2003).

42. W. Chen, J. M. Smeekens, R. Wu, Systematic study of the dynamics and half-lives of newly synthesized proteins in human cells. Chemical science 7, 1393–1400 (2016).

43. T. Mathieson et al., Systematic analysis of protein turnover in primary cells. Nature communications 9, 1–10 (2018).

44. A. M. Cunningham et al., Sperm transcriptional state associated with paternal transmission of stress phenotypes. The Journal of Neuroscience, JN-RM-3192-3120 (2021).

45. W. Gilbert, Origin of life: The RNA world. nature 319, 618–618 (1986).

46. L. E. Orgel, The origin of life on the earth. Scientific American 271, 76–83 (1994).

47. R. S. Moore et al., The role of the Cer1 transposon in horizontal transfer of transgenerational memory. Cell 184, 4697–4712. e4618 (2021).

48. J. V. McConnell, Memory transfer through cannibalism in planarians. (1965).

49. A. Bédécarrats, S. Chen, K. Pearce, D. Cai, D. L. Glanzman, RNA from Trained Aplysia Can Induce an Epigenetic Engram for Long-Term Sensitization in Untrained Aplysia. eNeuro, (2018).

50. P. Rajasethupathy et al., A role for neuronal piRNAs in the epigenetic control of memory-related synaptic plasticity. Cell 149, 693–707 (2012).

51. P. Spadaro, T. W. Bredy, Emerging role of non-coding RNA in neural plasticity, cognitive function, and neuropsychiatric disorders. Frontiers in genetics 3, 132 (2012).

52. C. M. Alberini, Transcription factors in long-term memory and synaptic plasticity. Physiological reviews 89, 121–145 (2009).

53. C. M. Alberini, M. Ghirardl, R. Metz, E. R. Kandel, C/EBP is an immediate-early gene required for the consolidation of long-term facilitation in Aplysia. Cell 76, 1099–1114 (1994).

54. K. C. Martin et al., Synapse-specific, long-term facilitation of Aplysia sensory to motor synapses: a function for local protein synthesis in memory storage. Cell 91, 927–938 (1997).

55. K. C. Martin et al., MAP kinase translocates into the nucleus of the presynaptic cell and is required for long-term facilitation in Aplysia. Neuron 18, 899–912 (1997).

56. S. Memczak et al., Circular RNAs are a large class of animal RNAs with regulatory potency. Nature 495, 333–338 (2013).

57. E. Lasda, R. Parker, Circular RNAs: diversity of form and function. Rna 20, 1829–1842 (2014).

58. W. Penfield, B. Milner, Memory deficit produced by bilateral lesions in the hippocampal zone. AMA Archives of Neurology & Psychiatry 79, 475–497 (1958).

59. L. R. Squire, The legacy of patient HM for neuroscience. Neuron 61, 6–9 (2009).

60. U. Dube et al., An atlas of cortical circular RNA expression in Alzheimer disease brains demonstrates clinical and pathological associations. Nature neuroscience 22, 1903–1912 (2019).

61. X. You et al., Neural circular RNAs are derived from synaptic genes and regulated by development and plasticity. Nature neuroscience 18, 603–610 (2015).

62. W. Chen, E. Schuman, Circular RNAs in brain and other tissues: a functional enigma. Trends in Neurosciences 39, 597–604 (2016).

63. I. P. Pavlov. (Oxford University Press, London. A PHYSIOLOGICAL MODEL OF PHOBIC ANXIETY A, 1927).

64. R. A. Rescorla, A. R. Wagner, A theory of Pavlovian conditioning. Classical Conditioning II: Current Theory and Research, (1971).

65. N. Mackintosh, An analysis of overshadowing and blocking. Quarterly Journal of Experimental Psychology 23, 118–125 (1971).

66. L. J. Kamin, in SYMP. ON PUNISHMENT. (1967).

67. R. R. Mowrer, S. B. Klein, Handbook of contemporary learning theories. (Psychology Press, 2000).

68. S. A. Josselyn, P. W. Frankland, Memory Allocation: Mechanisms and Function. Annual review of neuroscience 41, 389–413 (2018).

69. F. B. Krasne, J. D. Cushman, M. S. Fanselow, A Bayesian context fear learning algorithm/automaton. Frontiers in behavioral neuroscience 9, 112 (2015).

70. M. E. Stanton, Multiple memory systems, development and conditioning. Behavioural brain research 110, 25–37 (2000).

71. N. J. Murawski, A. Asok, Understanding the contributions of visual stimuli to contextual fear conditioning: a proof-of-concept study using LCD screens. Neuroscience letters 637, 80–84 (2017).

72. R. Bourtchouladze et al., Different training procedures recruit either one or two critical periods for contextual memory consolidation, each of which requires protein synthesis and PKA. Learning & Memory 5, 365–374 (1998).

73. E. R. Kandel, J. H. Schwartz, Molecular biology of learning: modulation of transmitter release. Science 218, 433–443 (1982).

74. V. Castellucci, H. Pinsker, I. Kupfermann, E. R. Kandel, Neuronal mechanisms of habituation and dishabituation of the gill-withdrawal reflex in Aplysia. Science 167, 1745–1748 (1970).

75. T. J. Carew, H. M. Pinsker, E. R. Kandel, Long-term habituation of a defensive withdrawal reflex in Aplysia. Science 175, 451–454 (1972).

76. W. Schreiber, A. Asok, S. Jablonski, J. Rosen, M. Stanton, Egr-1 mRNA expression patterns in the prefrontal cortex, hippocampus, and amygdala during variants of contextual fear conditioning in adolescent rats. Brain research 1576, 63–72 (2014).

77. A. Asok, L. W. Ayers, B. Awoyemi, J. Schulkin, J. B. Rosen, Immediate early gene and neuropeptide expression following exposure to the predator odor 2, 5-dihydro-2, 4, 5-trimethylthiazoline (TMT). Behavioural brain research 248, 85–93 (2013).

78. L. Luo, Architectures of neuronal circuits. Science 373, eabg7285 (2021).

79. A. Asok et al., A multisensory circuit for gating intense aversive experiences. bioRxiv,2021.2005.2001.441648 (2021).

80. D. M. Nielsen, L. S. Crnic, Automated analysis of foot-shock sensitivity and concurrent freezing behavior in mice. Journal of neuroscience methods 115, 199–209 (2002).

81. J. H. Choi et al., Interregional synaptic maps among engram cells underlie memory formation. Science 360, 430–435 (2018).

82. W. Wu, P. Ji, F. Zhao, CircAtlas: an integrated resource of one million highly accurate circular RNAs from 1070 vertebrate transcriptomes. Genome biology 21, 1–14 (2020).

83. S. R. Kassabov et al., A single Aplysia neurotrophin mediates synaptic facilitation via differentially processed isoforms. Cell reports 3, 1213–1227 (2013).

84. J. D. Sweatt, E. R. Kandel, Persistent and transcriptionally-dependent increase in protein phosphorylation in long-term facilitation of Aplysia sensory neurons. Nature 339, 51–54 (1989).

85. Y.-Y. Huang, K. C. Martin, E. R. Kandel, Both protein kinase A and mitogen-activated protein kinase are required in the amygdala for the macromolecular synthesis-dependent late phase of long-term potentiation. Journal of Neuroscience 20, 6317–6325 (2000).

86. M.dos Santos Corrêa et al., Relationship between footshock intensity, post-training corticosterone release and contextual fear memory specificity over time. Psychoneuroendocrinology 110, 104447 (2019).

87. M. Davis, D. I. Astrachan, Conditioned fear and startle magnitude: effects of different footshock or backshock intensities used in training. Journal of Experimental Psychology: Animal Behavior Processes 4, 95 (1978).

88. G. L. Quirarte, R. Galvez, B. Roozendaal, J. L. McGaugh, Norepinephrine release in the amygdala in response to footshock and opioid peptidergic drugs. Brain research 808, 134–140 (1998).

89. A. K. Rajbhandari, S. T. Gonzalez, M. S. Fanselow, Stress-enhanced fear learning, a robust rodent model of post-traumatic stress disorder. JoVE (Journal of Visualized Experiments), e58306 (2018).

90. D. J. Cai et al., A shared neural ensemble links distinct contextual memories encoded close in time. Nature 534, 115–118 (2016).

91. P. Namburi, R. Al-Hasani, G. G. Calhoon, M. R. Bruchas, K. M. Tye, Architectural representation of valence in the limbic system. Neuropsychopharmacology 41, 1697–1715 (2016).

92. W. B. Scoville, B. Milner, Loss of recent memory after bilateral hippocampal lesions. Journal of Neurology, Neurosurgery, and Psychiatry 20, 11–21 (1957).

93. S. Tonegawa, M. D. Morrissey, T. Kitamura, The role of engram cells in the systems consolidation of memory. Nature Reviews Neuroscience 19, 485–498 (2018).

94. N.-N. Zhang et al., A thermostable mRNA vaccine against COVID-19. Cell 182, 1271–1283. e1216 (2020).

